# Are infraslow oscillations the missing link between sleep and Alzheimer’s?

**DOI:** 10.64898/2026.04.09.717425

**Authors:** Demetrio Grollero, Victoria Gabb, Jonathan Blackman, Luisa de Vivo, Elisabeth Coulthard, Michele Bellesi

## Abstract

**INTRODUCTION:** Locus coeruleus and glymphatic dysfunction are linked both to Alzheimer’s disease (AD) and, recently, to infraslow oscillation in sleep spindle (sigma) activity (ISO). Here we hypothesise ISO integrity is a critical link between sleep and AD.

**METHODS:** We analyzed non-rapid eye movement sleep EEG from AD and controls, extracting ISO peak amplitude, intrinsic frequency, and bandwidth from the sigma-power time course. We assessed group differences and correlations with plasma biomarkers (Aβ42/40, pTau181 and 217, NfL, GFAP).

**RESULTS:** ISO peak amplitude was significantly reduced in AD, while intrinsic frequency and bandwidth were preserved. ISO peak amplitude positively correlated with Aβ42/40 ratio, and ISO bandwidth correlated with GFAP and NfL levels, and with lower verbal memory retention.

**DISCUSSION:** Such selective weakening of ISO in AD is consistent with LC dysfunction and impaired glymphatic cycling. ISO may be a novel mechanism and electrophysiological marker linking sleep microarchitecture to AD pathology.

## INTRODUCTION

Sleep architecture is profoundly altered in Alzheimer’s disease (AD), and growing evidence suggests that these disruptions are not merely symptomatic but mechanistically linked to neurodegenerative processes [1–3]. In particular, non-rapid eye movement (NREM) sleep, during which coordinated oscillatory activity supports memory consolidation, synaptic homeostasis, and glymphatic clearance, markedly deteriorates early in the disease course [4–6]. Among the key electrophysiological markers of NREM sleep, spindle-related sigma (12–16 Hz) activity has emerged as a sensitive indicator of thalamocortical circuit integrity and cognitive function [7,8]. Recent work has revealed that NREM sigma power is not constant but manifests infraslow oscillation (ISO), such that sleep spindles temporally cluster at <0.1 Hz [9,10].

A growing body of evidence indicates that these ISO dynamics are tightly coupled to oscillatory activity in the locus coeruleus (LC) and brain release of noradrenaline (NA) [10–12]. The LC exhibits rhythmic fluctuations in firing at ISO timescales, which produce corresponding modulations of NA release resulting in fluctuation of cortical arousal and sleep spindle expression during NREM sleep.

Furthermore, these oscillatory fluctuations in noradrenergic activity during NREM sleep modulate pericyte contraction, astrocytic end-feet geometry, and vascular tone, thereby promoting cerebrospinal fluid (CSF) influx and metabolic waste clearance, including β-amyloid [13,14].

The LC is among the earliest sites of tau-related pathology in AD, and impaired glymphatic function has been proposed as a key contributor to β-amyloid accumulation [15–17]. Thus, disruptions in LC-driven infraslow rhythms may mechanistically link sleep disturbances to the pathological protein deposition characteristic of AD.

In this study, we investigated the integrity of the ISO of sigma power in individuals with AD compared with age-matched healthy control participants. Here, we test whether breakdown of ISO coordination is evident in people affected by AD, reflecting LC dysfunction and impaired glymphatic cycling. We hypothesised that the integrity of the ISO of sigma power in individuals with AD would be reduced compared with age-matched healthy control participants.

## MATERIALS AND METHODS

### Participants

Thirty older adults (age > 50 years) were selected from the RESTED-AD cohort [18], a longitudinal observational study of sleep physiology in individuals with and without AD pathology. The sample included ten participants with AD and twenty cognitively healthy, age-matched controls. All participants met the RESTED-AD inclusion criteria and provided written informed consent. Details on recruitment, ethics, diagnostic procedures, and multimodal phenotyping have been described previously [18].

### Protocol and procedures

RESTED-AD is a naturalistic, home-based study designed to capture variability in sleep and behavior under real-world conditions. Following baseline clinical and neuropsychological assessment, participants underwent an intensive monitoring phase. Sleep was recorded for seven consecutive nights at home using a wireless multichannel portable EEG device (Dreem-2), with recordings self-initiated at habitual bedtime.

Cognitive performance was assessed using a word list memory task including immediate and next-morning recall after evening learning session. Blood samples were collected according to standardized procedures for quantification of plasma biomarkers, including Aβ42 and Aβ40, phosphorylated tau species (pTau-181 and pTau-217), neurofilament light chain (NfL), and glial fibrillary acidic protein (GFAP).

### EEG preprocessing and selection of NREM2 segments

All EEG preprocessing and analyses were performed in MATLAB using custom scripts together with FieldTrip and EEGLAB (2021). EEG data were band-pass filtered between 0.5 and 35 Hz using zero-phase FIR filtering. Artifacts were identified using an automated epoch-based rejection procedure based on kurtosis and joint probability. Contiguous artifact-free stage 2 NREM (NREM2) segments of at least 5 minutes were selected, and fixed-length 5-minute epochs were extracted for analysis. A frontal– central signal of interest was obtained by averaging the two available bipolar derivations, approximating centro-parietal regions where sigma activity is maximal.

### Sleep staging and quality control

Sleep staging was derived from automated hypnogram files generated by the portable EEG system. The Dreem algorithm, validated against expert-scored polysomnography, shows good agreement for sleep stage classification (∼83–84% accuracy) [19]. To minimize misclassification, all artifact-free NREM2 segments were visually inspected using a custom interface. Segments with residual artifacts or stage-inconsistent EEG features were excluded. Only NREM2 segments independently validated by two expert scorers were retained for analysis, ensuring reliable infraslow measurements.

### Data post processing and analyses

#### Isolation of ISO

ISO dynamics were characterized following established procedures described in [9] and extended in recent human studies [20]. Continuous sigma-band amplitude time courses were derived using time-frequency decomposition, and their power spectra were analyzed in the infraslow range (<0.1 Hz) to quantify rhythmic modulation of sigma activity. Sigma-band activity was estimated using Morlet wavelets (4-cycle wavelets over frequencies from 10 to 16 Hz in 0.2 Hz steps). The amplitude envelope was obtained by computing the absolute value of the complex Fourier coefficients and averaging it across frequencies at 0.1 second steps (10 Hz sampling). Time courses were detrended, demeaned, and smoothed using a moving-average filter (5 s window) to reduce activity unrelated to infraslow dynamics. Power spectral density of the sigma envelope was then computed using Welch’s method (window type:Hanning, 240s with 90% overlap), retaining spectral estimates in the 0–0.12 Hz range. To facilitate comparisons across nights and individuals, spectra were expressed in relative units by normalizing to median infraslow power and averaged across accepted segments within each night.

### ISO feature extraction and quality control

ISO features were extracted from relative power spectra using a model-based fitting approach. For each spectrum, a three-parameter (frequency, amplitude, width) Gaussian function with an additive constant term (constant baseline) was fitted. This formulation explicitly models the infraslow spectral peak as a localized oscillatory component superimposed on a non-oscillatory spectral background. Although spectra were expressed in relative units to account for variability in overall signal amplitude, retaining a baseline term allows residual broadband structure in the infra-slow range to be estimated directly from the data. This method avoids a priori assumptions about baseline shape (e.g., flat or predefined offsets) and accounts for aperiodic (1/f-like) components of EEG power spectra [21,22]. Fits were restricted to the 0.01–0.04 Hz range, previously shown to capture dominant ISO modulation of sigma power [9]. ISO peak frequency, peak amplitude, and bandwidth were defined from the Gaussian fit as the center frequency, height above baseline, and twice the standard deviation of the oscillatory component, respectively.

To ensure robust detection of meaningful ISO components, spectra were required to exhibit a clear spectral peak exceeding background variability and to show improved fit relative to a constant model. Spectra failing these criteria were excluded from further analyses.

### Power analysis

EEG spectral power was quantified during NREM2 and NREM3 using pre-processed data segments. Continuous NREM data were divided into non-overlapping 5-s epochs, and power spectra were estimated using a multitaper FFT approach over the 0.5-35 Hz range (using FieldTrip with a multitaper FFT approach with spectral smoothing of 2 Hz, zero-padding to the epoch length, and linear detrending per epoch; frequency range 0.5-35 Hz). Band-limited power was computed for delta (0.5–4 Hz; SWA), alpha (8–12 Hz), sigma (12–16 Hz), and beta (18–35 Hz). Total broadband power was defined as the integrated power over 0.5-35 Hz. Relative power was expressed as the percentage contribution of each band to total broadband power. Epoch-level power estimates were averaged within each sleep stage and night, and subsequently across nights to obtain subject-level mean power values for NREM2 and NREM3. Relative power measures were interpreted as reflecting the spectral composition of NREM activity rather than absolute signal amplitude.

### Memory evaluation

Memory performance was quantified using three measures: recognition sensitivity (d′), memory retention, and effective recall. Recognition sensitivity (d′) was computed according to signal detection theory using true positives, false negatives, false positives, and true negatives, providing a measure of the ability to discriminate previously learned items from novel ones. Memory retention was defined as the ratio of delayed to immediate free recall, indexing overnight preservation of learned material. Effective recall was calculated as the number of correctly recalled items minus intrusions during immediate recall. Standard corrections were applied to ensure numerical stability of the derived measures. Participants with missing values were excluded from analyses.

## Statistical analysis

Statistical analysis was performed using MATLAB and GraphPrism (10.6.01). Group differences in ISO-features were evaluated at the night level. Between-group differences were assessed using parametric statistical tests, given normal data distribution after log-transformation. For correlations with plasma biomarkers and memory measures, ISO features were further averaged across nights to obtain subject-level estimates. All statistical tests were two-tailed with a significance threshold of α = 0.05.

## RESULTS

### Dataset quality description

After quality control, one participant was excluded due to poor EEG data quality. A total of ∼2,050 artefact-free 5-minute NREM2 segments (∼170 h of NREM sleep) were retained across 152 nights from 29 participants. Nights with fewer than two valid segments (10.53%) or without a dominant infra-slow oscillatory peak (7.9%, see methods section) were excluded, resulting in a final sample of 124 nights (AD: 41 nights from 9 participants; controls: 83 nights from 18 participants). No differences were observed between groups in the number of nights included per subject (P = 0.95) or in the average number of valid NREM2 segments per night (P = 0.21; Supplementary Figure 1).

### ISO of the sigma power is reduced in AD

We quantified the modulation of ISO by extracting three features from each participant’s ISO spectrum of the sigma power time course: (i) the ISO peak amplitude, reflecting the strength of the rhythmic oscillation; (ii) the ISO intrinsic frequency, representing the dominant oscillatory rate within the infraslow range; and (iii) the ISO bandwidth, capturing the spread of spectral power around the peak frequency. Across groups, a clear reduction emerged in the ISO peak amplitude in AD patients compared with healthy controls (-28,03 ± 6.88 %, P = 0.003, Figure 1), indicating a marked weakening of the infraslow oscillation amplitude of sigma activity. In contrast, neither the ISO intrinsic frequency (P = 0.77) nor the ISO bandwidth (P = 0.79) differed significantly between groups (Figure 1), suggesting that while the rhythmic structure and frequency of the oscillation are preserved, its overall strength is selectively diminished in AD. In addition, broad EEG spectrum analysis did not reveal significant reductions in delta or spindle power in AD relative to controls (Supplementary Figure 2), as observed in prior studies [23]. Moreover, none of ISO features correlated with the extent of delta in patients and controls (Amplitude: r = -0.06, P = 0.8; Frequency: r = -0.04, P = 0.85; Bandwidth: r = 0.18, P = 0.4).

**Figure 1.**
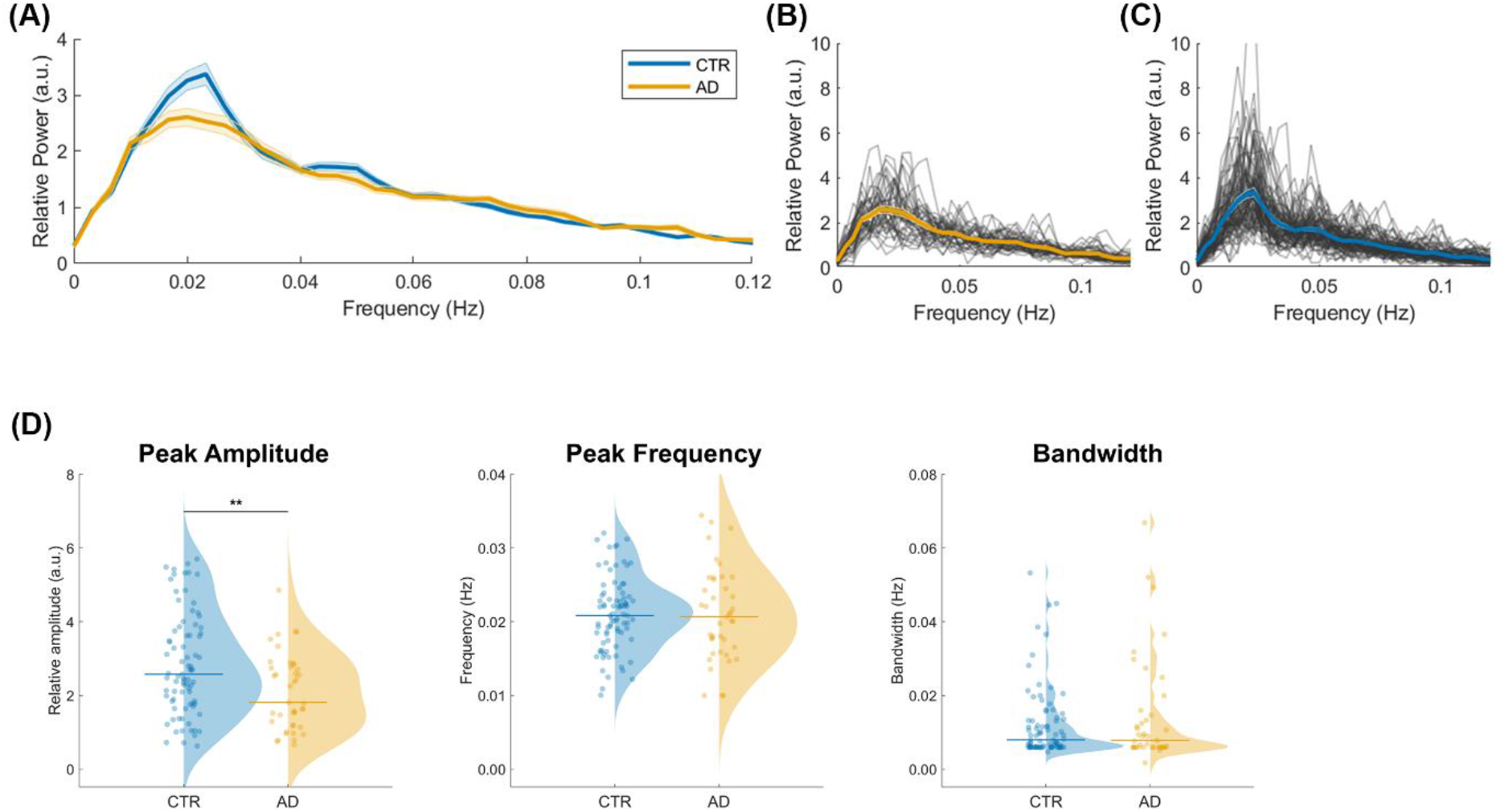
Group differences in ISO spectra and features. **A-C**. Group-average night-level spectra for CTR (blue) and AD (yellow) are shown as mean ± SEM across nights. Individual night-level spectra (thin grey lines) for AD (**B**) and CTR (**C**) are also presented, with the corresponding group means ± SEM overlaid. Spectra are expressed in relative units (a.u.) with respect to the 0.006-0.1 Hz frequency band. For visualization only, ultra-low-frequency peaks (<0.006 Hz), attributable to residual baseline offsets and outside the predefined infra-slow range, were masked prior to averaging. **D**. Distribution of ISO features in CTR (blue) and AD (yellow). Panels display peak amplitude (left), peak frequency (centre), and bandwidth (right) of the infra-slow modulation of sigma power. Individual dots represent single nights, half-violins depict kernel density estimates, and horizontal lines indicate the median value within each group. Between-group differences were assessed using t-tests; **p < 0.01.

### ISO features correlate with plasma biomarkers of disease

We next assessed whether alterations in infraslow oscillation features were related to established plasma biomarkers of AD pathology and neurodegeneration. A significant positive correlation emerged between ISO peak amplitude and the Aβ42/40 ratio (Table 1), such that individuals with greater infraslow oscillation tended to exhibit more favorable amyloid profiles. Notably, AD patients clustered at the lower end of both ISO peak amplitude and Aβ42/40 ratio (Figure 2A), consistent with a coupling between diminished infraslow oscillation and greater amyloid burden.

**Table 1.** ISO Features correlations with plasma biomarkers.

| ISO Feature | Aβ42/40) | pTau 181 | pTau 217 | GFAP | NFL |
| --- | --- | --- | --- | --- | --- |
| Amplitude | R=0.5, * **P=0.02** | R=0.2, P=0.37 | R=0.3, P=0.16 | R=0.15, P=0.5 | R=0.4, P=0.1 |
| Frequency | R=-0.01, P=0.97 | R=0.09, P=0.7 | R=0.1, P=0.6 | R=-0.3, P=0.16 | R=-0.3, P=0.13 |
| Bandwidth | R=-0.04, P=0.8 | R=0.4, P=0.09 | R=0.4, P=0.07 | R=0.5, * **P=0.03** | R=0.8, * **P<0.001** |

**Figure 2.**
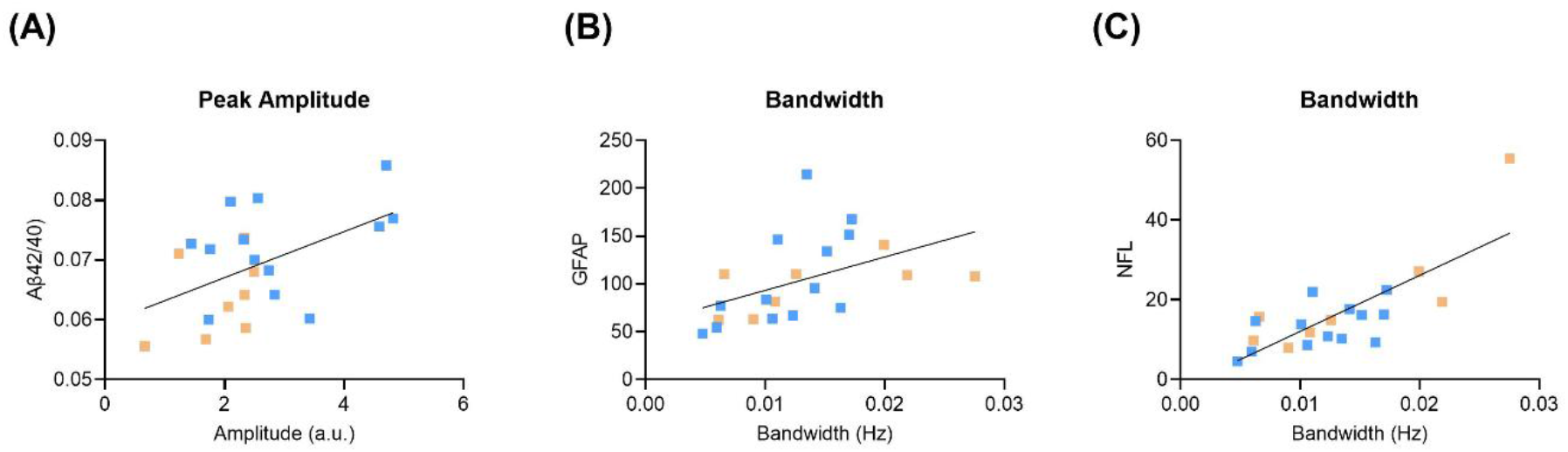
Significant ISO features - plasma biomarkers correlations. **A**. Correlation between ISO Peak Amplitude and Aß42/40 ratio. **B-C**. Correlation between ISO Bandwidth with GFAP (**B**) and NFL (**C**) concentrations. For all panels, yellow squares represent AD, while the blue ones are CTR.

In addition, we observed that the ISO bandwidth showed positive correlations with plasma levels of GFAP and NfL (across AD and control groups), markers of astroglial activation and axonal injury, respectively (Table 1). Participants with broader infraslow spectral bandwidth tended to present with higher concentrations of these biomarkers (Figure 2B-C), suggesting that more heterogeneity in spectral features of infraslow activity may reflect underlying glial and neurodegenerative processes. No significant associations were found for the ISO intrinsic frequency (Table 1).

### ISO bandwidth is associated with memory impairment

Finally, we examined the relationship between ISO features and memory performance on a word list task. ISO bandwidth showed a significant negative correlation with word retention (R = −0.49, P = 0.03), indicating that individuals with broader infraslow bandwidth exhibited poorer memory performance (Table 2). Other ISO features were not significantly related to any of the memory task parameters (Table 2).

**Table 2.** ISO Features correlations with memory metrics.

| ISO Feature | dprime | Retention | Recall |
| --- | --- | --- | --- |
| Amplitude | R=0.1, P=0.65 | R=0.16, P=0.5 | R=0.3, P=0.2 |
| Frequency | R=-0.04, P=0.85 | R=0.09, P=0.7 | R=0.2, P=0.3 |
| Bandwidth | R=-0.43, P=0.06 | R=-0.5, * **P=0.03** | R=-0.24, P=0.3 |

## DISCUSSION

This study examines whether alterations in ISO dynamics, previously proposed to reflect LC activity and influence glymphatic function, are detectable in human sleep and relate to molecular markers of AD pathology. By quantifying ISO features from overnight EEG, we demonstrate a selective reduction in ISO peak amplitude in people with clinically-diagnosed AD, while intrinsic frequency and bandwidth remain preserved. This pattern reveals a specific weakening of the rhythmic modulation that structures sigma activity, suggesting impaired LC-driven arousal cycling during NREM sleep.

Our findings offer a novel perspective by showing that ISO alterations are not only present in AD but are meaningfully related to plasma biomarkers of amyloid dysregulation and neurodegeneration. The positive association between ISO amplitude and Aβ42/40 ratio indicates that preserved infraslow modulation accompanies a more favorable amyloid profile, while correlations between ISO bandwidth and GFAP and NfL suggest that additional spectral properties track glial activation and axonal injury. Specifically, broader ISO bandwidth corresponds to a less sharply defined infraslow oscillatory peak and greater dispersion of power across frequencies, a pattern consistent with reduced ISO stability. No significant associations were found for the ISO intrinsic frequency. In parallel, the association between broader ISO bandwidth and poorer word retention provides initial evidence that alterations in infraslow sigma modulation may also relate to sleep-dependent memory processes impaired in AD [7,24]. Finally, ISO features did not correlate with delta power, the canonical electrophysiological marker of sleep depth frequently altered in AD [23,25]. This dissociation suggests that ISO abnormalities are not driven by reduced sleep intensity but instead reflect a distinct regulatory process, consistent with recent observations showing that NREM noradrenergic oscillations are more tightly coupled to light sleep features, such as microarousals, than to slow-wave sleep per se [13].

Several limitations should be acknowledged. The study is cross-sectional, preventing direct inference about temporal ordering between ISO disruption and biomarker changes. LC function and glymphatic activity were not measured directly, and EEG-derived ISO features represent downstream proxies rather than mechanistic readouts. Sample size and technical factors with recording device (such as bipolar derivations or non-standard electrode positions) may limit detection of delta or spindle power and subtler associations.

Despite these limitations, the present findings have clear implications. They suggest that reduced ISO amplitude may reflect a dysfunction of LC-glymphatic regulatory systems, offering a testable framework for understanding how sleep microarchitecture contributes to AD progression. Specific next steps will include: (1) longitudinal evaluation to determine whether ISO amplitude predicts cognitive decline or biomarker trajectories; (2) integration with LC imaging (neuromelanin MRI or noradrenergic PET) to directly link ISO alterations to LC integrity; and (3) intervention studies targeting noradrenergic tone or arousal stability to assess whether ISO amplitude can be restored and whether such restoration enhances glymphatic function or reduces pathological protein burden.

In conclusion, this work identifies weakened infraslow modulation of sigma activity as a potential electrophysiological signature of AD-related network and clearance dysfunction. Establishing ISO features as accessible, biologically grounded biomarkers may open new avenues for early detection and mechanistically guided therapeutic strategies.

## AUTHOR CONTRIBUTIONS

Conceptualization: MB. Data analysis: DG, MB. Data curation and graphics: DG. writing—original draft: DG, MB; writing—review and editing: all authors.; project supervision and funding: MB, LdV, EC.

## ACKNOWLEDGMENTS

The authors sincerely thank the Remote Evaluation of Sleep to Enhance Understanding of Early Dementia (RESTED) study participants and their relatives and partners who supported them to take part. They would also like to thank the ReMemBr group lived experience experts for their contributions to improving the study design and supporting them to make the study as accessible as possible. They would also like to thank the North Bristol NHS Trust Respiratory Physiology department and the Bristol Bioresource Laboratories, Anglia Ruskin University Biomarker Lab, and UK DRI Fluid Biomarker Laboratory for providing their services and support for our saliva and blood biomarker analyses.

Finally, the authors would like to thank the teams at Dreem Research, Cognitron, Join Dementia Research, and Ewa G Truchanowicz and Phil Reay, and their teams at Dignio, United Kingdom. We also thank Davide Marzoli for helpful methodological discussions.

The RESTED study was funded by charitable organizations (BRACE, Alzheimer’s Research UK, the Bristol and Weston Hospitals Charity [previously Above and Beyond], and the David Telling Charitable Trust), the NIHR Bristol Biomedical Research Centre, and philanthropic donations from S Scobie and A Graham. JB also received funding from Alzheimer’s Research United Kingdom (ARUK) supported by the Margaret Jost Fellowship and the Don Thoburn Memorial Scholarship. MB was supported by ARUK (PPG2020A-023) and LdV by the Giovanni Armenise-Harvard Foundation (CDA).

The funders had no role in the trial design, patient recruitment, data collection, analysis, interpretation, writing of the manuscript, or the decision to submit it for publication.

## CONSENT STATEMENT

All participants provided written informed consent, and the study was approved.by the Health Research Authority (Yorkshire and the Humber-Bradford Leeds Research Ethics Committee, reference 21/YH/0177).

## DATA AVAILABILITY STATEMENT

Data and informed consent forms are available upon request.

## SUPPLEMENTARY MATERIAL

**Supplementary Figure 1.**
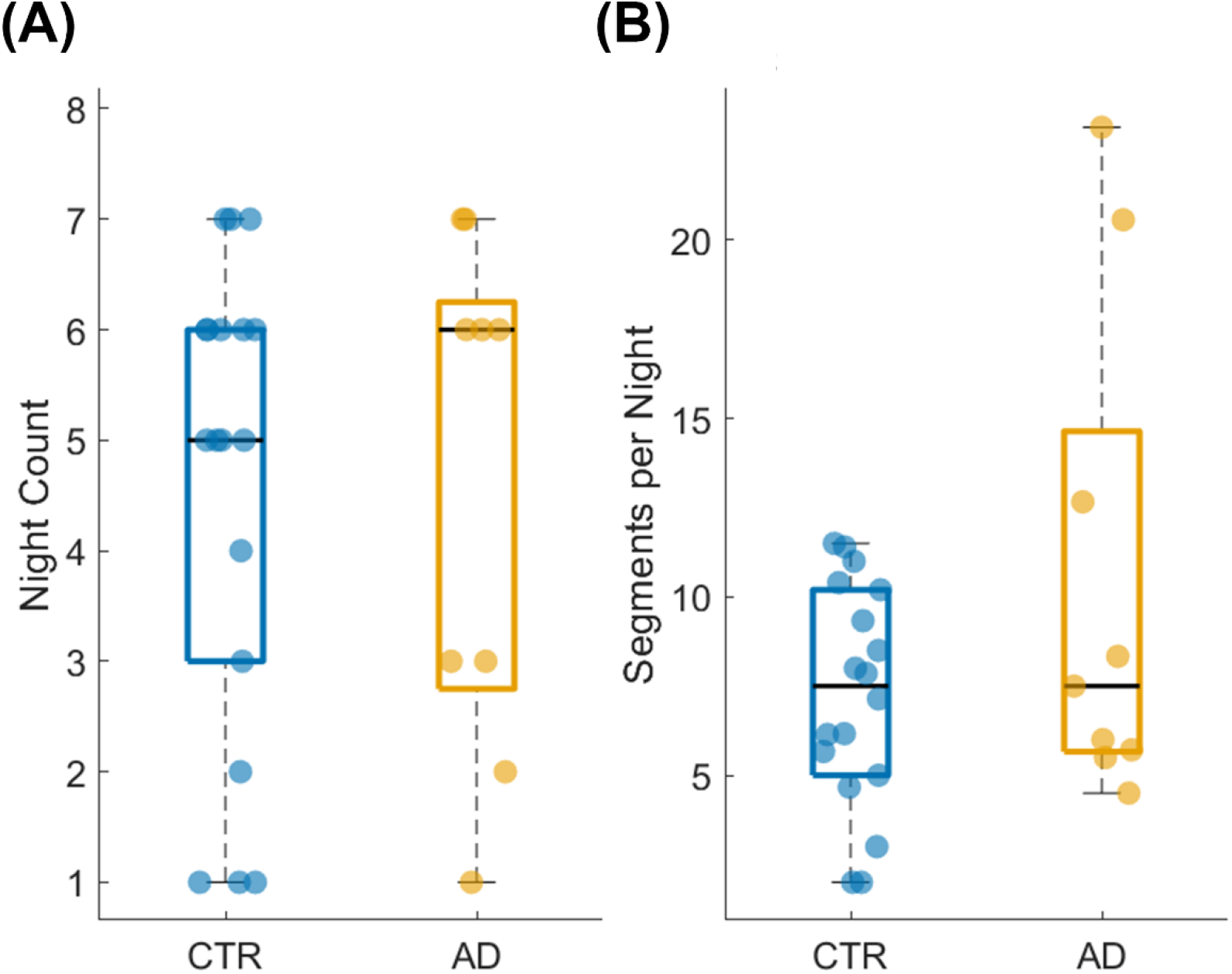
Group-wise data availability for ISO analyses. **A**. Distribution of the number of nights retained per subject in CTR and AD groups after preprocessing and quality control. **B**. Differences of the average number of valid NREM2 segments per night for each subject. Boxes indicate the interquartile range, central lines the median, whiskers the data range, and dots individual subjects. No significant group differences were observed in the number of nights retained or in the average number of segments per night (all *p* > 0.2).

**Supplementary Figure 2.**
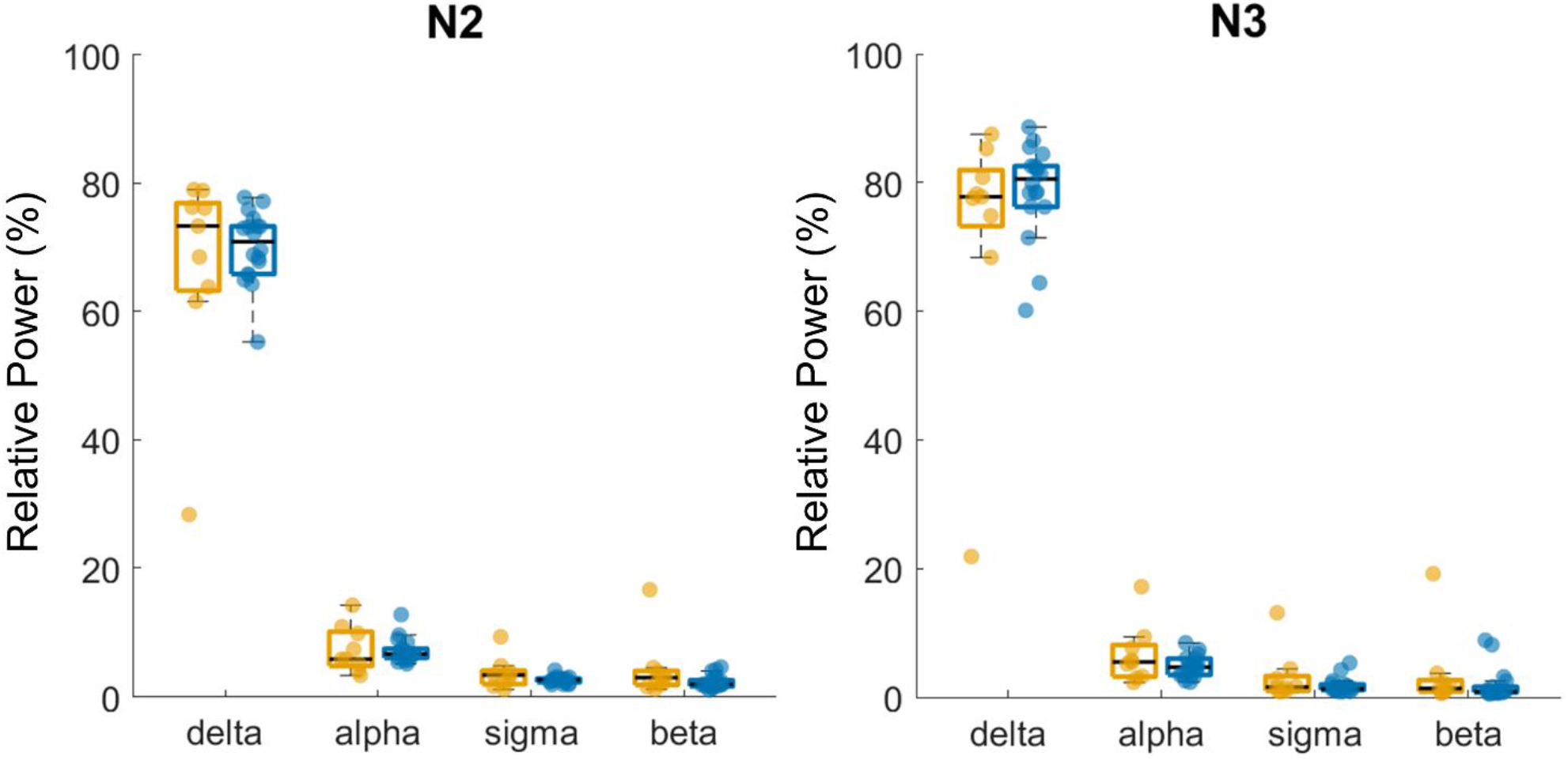
Average power for common frequency bands in AD and CTR in N2 and N3 REM sleep. Spectral power was computed from the bipolar fronto–occipital derivation within NREM2 and NREM3 time periods. Power spectra were estimated using a multitaper FFT approach. Relative band power was expressed as a percentage of total power.

